# Quantitative phosphoproteomics uncovers the signalling dynamics of hallucinogenic psychedelics

**DOI:** 10.1101/2025.11.24.690190

**Authors:** Sandra M. Martin-Guerrero, Marco Taddei-Tardon, Jessica L. Maltman, Sritanvi Potukanuma, Javier González-Maeso, Pedro R. Cutillas, Juan F. Lopez-Gimenez

## Abstract

Psychedelic drugs can induce intense changes in perception and thought, and some also promote long-lasting adaptations in brain circuits that are being explored for treatment of mood and anxiety disorders. How these compounds differ at the level of intracellular signalling, and how hallucinogenic drugs diverge from related non-hallucinogenic forms, is poorly understood. A central question is whether a shared molecular fingerprint distinguishes hallucinogenic psychedelic action from other forms of receptor activation in neurons. Here, we show that chemically diverse psychedelics trigger a coordinated reorganisation of phosphorylation patterns across many proteins in neural cells, and that this global signalling response contains a distinct signature that separates hallucinogenic compounds from non-hallucinogenic counterparts of similar structure. We use a glycolysis-regulating transcription factor as an example of the signature’s functional relevance to show that hallucinogenic psychedelics, but not their non-hallucinogenic analogues, enhance markers of glycolytic metabolism. These findings reveal that hallucinogenic and non-hallucinogenic psychedelics engage separable intracellular architectures, and establish a framework for understanding how different psychoactive compounds couple receptor activation to specific cellular states. More broadly, this work opens a path to using signalling fingerprints to guide the design of psychedelic-inspired therapeutics with tailored behavioural and metabolic profiles.

---

Over the past decade there has been renewed interest in using psychedelic drugs to treat neuropsychiatric disorders that do not respond to standard therapies, supported by preclinical evidence that these compounds induce lasting structural and functional plasticity in prefrontal cortical circuits^1^. At the same time, psychedelics produce acute and profound alterations of consciousness, raising the unresolved question of whether their enduring neuroplastic effects can be dissociated from transient hallucinogenic states^2–4^. Psychedelic-like activity can be produced by chemically diverse agents, including serotonergic tryptamines such as psilocybin and dimethyltryptamine (DMT), ergolines such as lysergic acid diethylamide (LSD), phenethylamines such as mescaline and 2,5-dimethoxy-4-iodoamphetamine (DOI), and the dissociative anaesthetic ketamine. Whereas serotonergic compounds act primarily as agonists at the 5-HT2A receptor^5,6^, a G-protein coupled receptor (GPCR), ketamine produces its effects predominantly through non-competitive blockade of NMDA receptors and the engagement of distinct downstream signalling adaptations^7^. Despite intensive study, the intracellular signalling circuits reorganized by these drugs, and how they relate to both hallucinogenic and plasticity-related outcomes, remain poorly defined^8^. A distinctive experimental advantage is the availability of closely related non-hallucinogenic analogues that retain the ability to promote structural and functional neuroplasticity while lacking hallucinogenic activity^9–11^, providing a framework to disentangle mechanisms of psychedelic-induced plasticity from those underlying acute alterations in perception.

To delineate signaling properties that discriminate hallucinogenic psychedelics from their non-hallucinogenic analogs, we profiled the phosphoproteomic landscape induced by a battery of such compounds on a previously described in vitro model based on neural cells. Importantly, because this model comprises neurons and glia differentiated from murine neural stem cells, it contains the signalling machinery engaged by G-protein-coupled receptor signalling in native nervous tissue ^12^ (Fig. 1a). Our serotonergic psychedelic panel includes psilocin, DMT, LSD, and DOI, as well as the non-hallucinogenic analogs Ariadne, 2BrLSD, and lisuride (Supplementary Fig. 1), and further included ketamine, a non-serotonergic hallucinogenic compound, and BDNF, an endogenous neurotrophin that binds to TrkB receptors and is implicated in neuroplasticity induced by psychedelics^13,14^, which served as a positive physiological control. To capture temporal dynamics of intracellular signaling, cells were treated for 5, 15, and 30 minutes, with each condition precisely time-matched to vehicle controls.

**Fig. 1.**
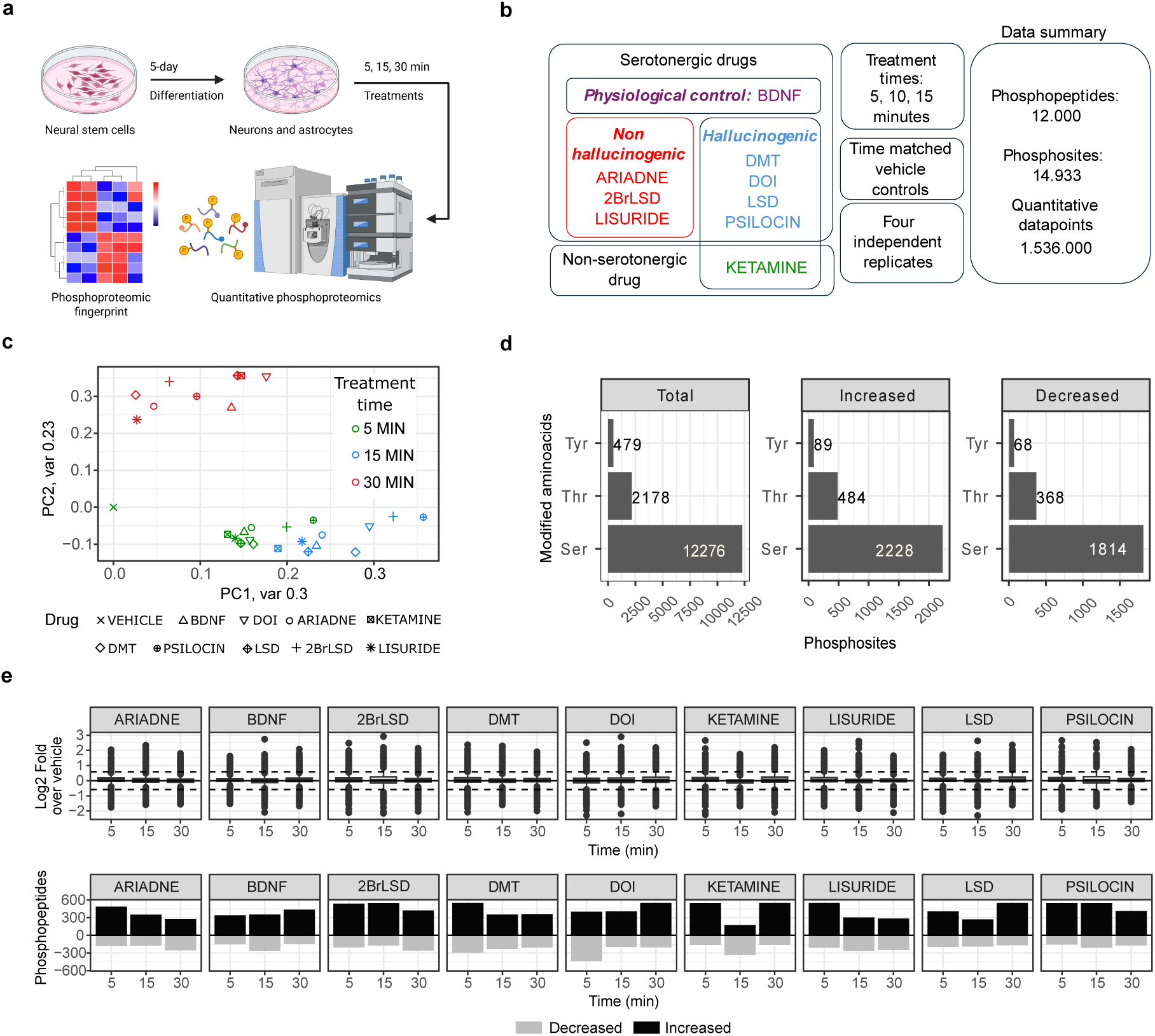
Phosphoproteomics of neural cells reveals extensive signaling changes induced by psychedelic compounds and BDNF. **a.** Experimental design. Neural stem cells were differentiated in media for 5 days into neurons and glia, subsequently treated with psychedelic compounds and BDNF at the time points shown and subjected to quantitative phosphoproteomics analysis. **b**. Experimental conditions used in this study (drugs used, treatment times and biological replicates) and data summary of phosphopeptides and phosphosites identified in the study. **c**. Principal component analysis (PC) using global phosphoproteomics data as input showing grouping of experimental conditions by treatment time. **d.** Number of phosphosites identified in the study grouped by modified amino acid (Ser, Thr or Tyr). **e.** Distribution of phosphopeptides Log2 Fold changes (treatment versus vehicle) (top plots) and number of phosphopeptides (bottom plots) found to be significantly modulated at the indicated times and drugs.

Quantitative phosphoproteomic analysis by LC–MS/MS followed by computational analysis identified 12,000 unique phosphopeptides encompassing 14,933 phosphorylation sites (Fig. 1b, Supplementary Information 1), distributed across serine (12,276), threonine (2,178), and tyrosine (479) residues (Fig. 1d). Experiments were conducted in four independent biological replicates, yielding a final phosphoproteomic dataset comprising more than 1.5 × 10⁶ quantitative data points (Supplementary Information 2). Following batch correction and averaging across replicates, unsupervised principal component analysis (PCA) of the complete dataset revealed clustering of samples primarily according to treatment duration, with phosphoproteomes from the 30-minute condition clearly segregating from those obtained after 5 and 15 minute treatments (Fig. 1c). Applying a fold-change threshold of 4.5 standard deviations from the mean, we identified 3,662 phosphorylation sites significantly regulated by at least one treatment, of which 2,801 were increased and 2,250 decreased (Fig. 1e). For each time point, individual drug treatments produced a greater number of upregulated than downregulated phosphorylation sites, with overall counts ranging from approximately 300 to 900 regulated sites per compound and time point (Fig. 1e). The total number of modulated sites does not correspond to the sum of increased and decreased events, as several sites exhibited bidirectional regulation across time points. Across treatments, the most recurrently regulated phosphosites mapped to protein kinases, G-protein–coupled receptors, small GTPases and their regulators, epigenetic modifiers and cytoskeletal proteins, with ontology analysis indicating enrichment for axonogenesis, neuronal migration, regulation of GTPase activity and other phosphorylation-dependent processes in neuronal projections and euchromatin (Extended Data Fig. 1 and Fig. 2).

**Fig. 2.**
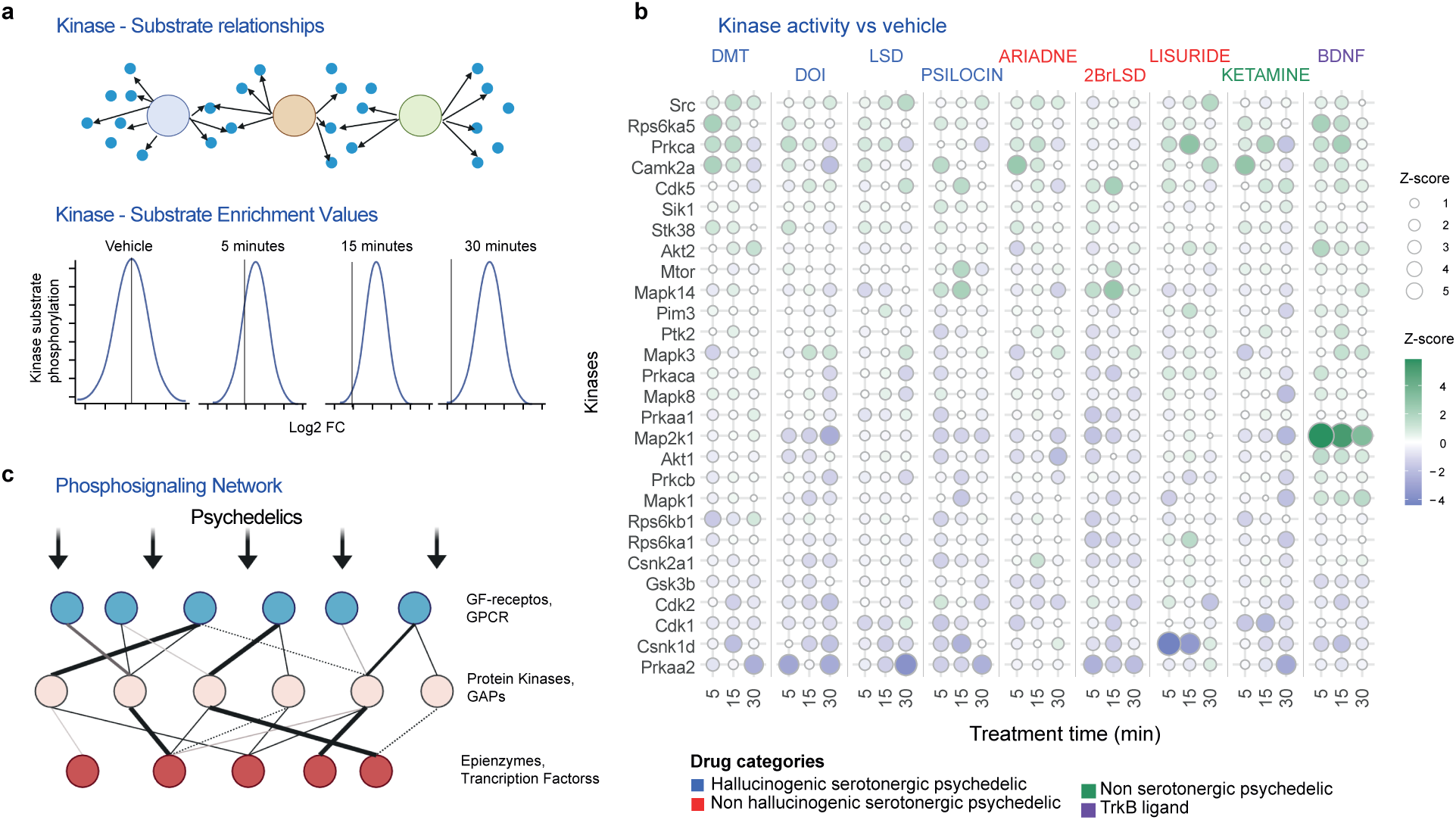
Computational analysis of phosphoproteomics data uncovers canonical and non- canonical signalling engaged by psychedelics drugs. **a.** Kinase substrate enrichment analysis (KSEA) links kinases to their substrates (top) and allows measuring kinase activation by comparing distribution of phosphorylation of substrates for given kinases relative to all phosphorylation sites (bottom). **b.** Results of KSEA depicting z- scores for the named kinases across the indicated time points and drugs. Colour and size of datapoints are proportional to kinase activities modulated by treatments. **c.** Approach to assemble regulatory phosphorylation networks linking cell surface receptors with protein kinases, epigenetic regulators and transcription factors.

Previous studies on functional selectivity mainly focused on 5-HT2A receptor have described how psychedelic agonists modulate the trafficking and activity of canonical intracellular pathways downstream of GPCRs and RTKs^15–17^. However, these studies have often produced inconclusive or contradictory results on the precise signaling cascades triggered downstream these receptors^18^. To obtain an unbiased assessment of canonical signaling events modulated by our treatments, we performed Kinase Substrate Enrichment Analysis^19^ (KSEA, Fig. 2) of the phosphoproteomics data (Supplementary Information 3). Most psychedelics reduced KSEA activity scores for kinases within the MAPK pathway, including MEK1 (Map2k1), ERK1 (Mapk3), and ERK2 (Mapk1), as well as for PKB (Akt1 and Akt2). As a positive control for neuronal plasticity processes, the TrkB receptor agonist BDNF increased the activity scores of these kinases, consistent with the established roles of MAPK and AKT signaling downstream of receptor tyrosine kinases in promoting neurotrophic responses^20^. In contrast, activity scores for Src, PKA, CaMK2A, and the stress-activated kinase p38 (Mapk14) increased following treatment with psychedelics and BDNF. Activity scores for mTOR also tended to increase, whereas those for several cell cycle regulators (CDK1, CDK2, CSNK1D) and AMPK (Prkaa2) decreased (Fig. 2b). These findings are consistent with the reciprocal regulation of mTOR and AMPK activities that maintains cellular metabolic homeostasis. Overall, our kinase activity data show that, contrary to prevailing assumptions in the field, psychedelics unexpectedly suppress MAPK and AKT signaling pathways while inducing changes in canonical signaling that only partially overlap with those elicited by BDNF-related neuronal plasticity mechanisms^13,14^. Consistent with the kinase-substrate enrichment analysis, examination of canonical pathway readouts showed that psychedelics and BDNF exert opposing effects on AKT and ERK activation while commonly enhancing mTOR- and focal adhesion kinase–linked signalling, validating the inferred pathway regulation (Extended Data Fig. 3).

We next aimed to investigate how psychedelics influence the propagation of downstream signaling cues on a broader scale. Treatments induced phosphorylation on multiple growth factor receptors (GFRs) and GPCRs, suggesting the engagement of diverse transactivation mechanisms (Extended Data Fig. 4). GFRs whose phosphorylation was affected by psychedelic treatment included the insulin receptor (INSR) at Y1179, the epidermal growth factor receptor (EGFR) at Y1110 and S1104, and the opioid growth factor receptor (OGFR) at S442 and S403. Similarly, GPCRs showing the most pronounced increases in phosphorylation included GPR161, ADGRL3, and GPRC5B, receptors previously described as regulators of neuronal morphology and synaptic connectivity^21–23^. Sites on protein kinases showed among the strongest phosphorylation changes following treatment, as indicated by robust increases at specific residues on MTOR, MAP3K1, MAP4K4, RIOK1, MARK1, and PRKCI. In addition, proteins involved in receptor signaling and epigenetic regulation were also affected^24^.

Phosphorylation changes were detected on enzymes from acetyltransferase, deacetylase, demethylase, and methyltransferase families (Extended Data Fig. 4b). Among acetyltransferases, NAA10, KAT5, NAA30, and KAT7 showed the most prominent increases in phosphorylation after treatment. The deacetylases HDAC1, HDAC4, HDAC5, HDAC9, and MIER1 also displayed marked phosphorylation increases. These data highlight the extensive signaling reorganization induced by psychedelics across proteins with diverse cellular functions.

To explore the relationships among phosphosites modulated by psychedelic treatments, we constructed a regulatory network in which phosphosites are represented as nodes, and the edges between them are weighted according to the temporal similarity of their fold change profiles across treatments, measured as correlation coefficients (Fig. 2c and Extended Data Fig. 5). This analytical approach is conceptually analogous to the construction of gene regulatory networks, where transcription factors and their targets are linked based on co-regulatory patterns. The resulting phosphosite network provides a framework for identifying coordinated phosphorylation events and for generating hypotheses regarding the propagation of signaling cascades downstream of receptor activation. For instance, the phosphorylation site on the GPCR ADGRB2 at S12169 showed strong associations with multiple sites on AKT isoforms, including AKT3 at T440, AKT2 at Y456, and AKT1 at T450, as well as with phosphorylation sites on the upstream kinase PDPK1 (S25) and downstream effectors RPS6KC1 (S587 and S527). Similarly, the GPCR FZD3 at S588 was associated with phosphorylation on kinases such as STK32C (S17), TNIK (S659), DCLK1 (T311), and MAP kinases involved in stress responses, including MAP4K4 (S621, S687, S683, and S691) and MAP3K3 (S340) (Extended Data Fig. 5a). Additional examples include associations between GPCRs and downstream GTP regulatory proteins (Extended Data Fig. 5b), as well as between protein kinases and epigenetic regulators (Extended Data Fig. 5c). Notably, phosphorylation on histone deacetylase HDAC1 at S421 and S423 was strongly associated with sites on kinases including SIK3 (a positive regulator of MTOR)^25^, AAK1 (involved in endocytosis)^26^, MAP3K1, ROCK2 (a key regulator of actin cytoskeletal dynamics)^27^, and LIMK2 (which acts downstream of ROCK2)^28^. Likewise, phosphorylation sites on methyltransferases MECOM (S1041), CMTR1 (S52), and MEPCES (S192) were associated with multiple kinases such as MAP4K3, CAMK2B, TNIK, and DCLK1. Together, these data provide a comprehensive resource for hypothesis generation regarding the interactions between signaling networks and epigenetic regulation, which may underlie long-term cellular effects elicited by acute psychedelic exposure.

As noted previously, this study included serotonergic hallucinogenic analogues that lack psychedelic activity and thus display head-twitch response (HTR) negative profiles—a well-established behavioral correlate of human hallucinogenic potential in mice^6,9,10,29^. Nevertheless, HTR assays were conducted to enable a direct comparison of the hallucinogenic potency of the compounds used in the phosphoproteomics experiments under identical experimental conditions. The results on this *in vivo* compound characterization (Extended Data Fig. 6) revealed marked differences in HTRs between the phenethylamines DOI and Ariadne and the ergolines LSD, 2BrLSD, and lisuride. Among the hallucinogenic compounds, DOI induced the most robust HTR, whereas DMT evoked the weakest response, consistent with previous observations for this tryptamine^29^. Time-course analysis further confirmed the absence of a significant HTR following administration of Ariadne, 2BrLSD, lisuride, ketamine, or DHF, the later serving as a systemically active TrkB agonist used as a substitute for BDNF (Extended Data Fig. 6a).

To further characterise these effects, differences in downstream signalling induced by psychedelics were examined based on their hallucinogenic properties by integrating the area under the curve of time response profiles for each phosphopeptide and compared these values across hallucinogenic compounds relative to their non-hallucinogenic counterparts (Extended Data Fig. 7) Using a tolerance of 4 units of standard deviation, we identified a phosphoproteomic signature consisting of 73 sites (Fig. 3a) (Supplementary Information 4), which allowed classification of phosphosites according to hallucinogenic potential (Fig. 3a and 3b). Ontology analysis revealed that sites in this signature of hallucinogenic potential were located on proteins involved in the spliceosome, actin filaments, and the centriole, with established roles in thalamus development, dendritic spine maintenance, and biological processes related to glucose homeostasis, glycolysis, and response to starvation (Fig. 3c and Extended Data Fig. 8), suggesting that these processes are involved in mediating the hallucinogenic properties of the tested compounds.

**Fig. 3.**
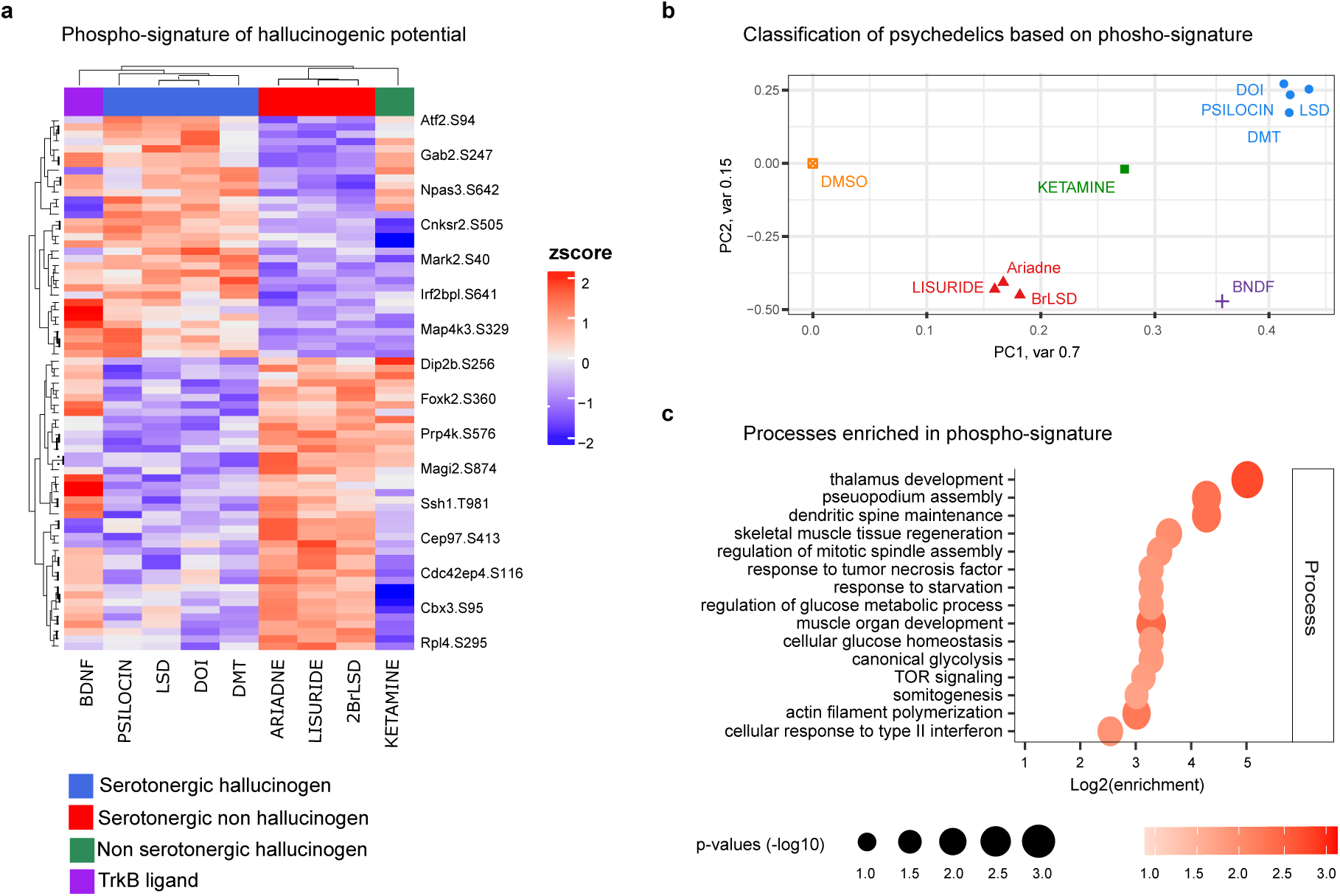
Identification of a phosphoproteomic signature that distinguishes non-hallucinogenic serotonergic psychedelics from hallucinogenic serotonergic psychedelics. **a.** Heatmap showing the differential phosphorylation of 73 phosphopeptides induced by the named drugs. **b.** Principal component analysis (PC) based in the signature obtained in **a**. **c.** Gene Ontology analysis of processes enriched in the phospho-signature of hallucinogenic versus non-hallucinogenic psychedelic activity. The color and size of the datapoints reflect the significance of each term’s enrichment. The x-axis shows the enrichment of each gene ontology term.

Of note, phosphorylation of FOXK2 at S360 was among the sites showing the largest differences between compound classes (Fig. 4a and 4b). FOXK2 is a transcription factor implicated in the regulation of glucose homeostasis and aerobic glycolysis^30^, consistent with the ontology analysis revealing an enrichment of these processes within the hallucinogenic signature (Fig. 3c). These observations led us to hypothesize that phosphorylation-dependent activation of FOXK2 may enhance glycolytic flux. To test this possibility, we performed lactate production assays to determine whether FOXK2 phosphorylation was associated with increased glycolytic activity. Treatment with hallucinogenic compounds elevated lactate levels by approximately 2- to 4.5-fold relative to vehicle, whereas their non-hallucinogenic counterparts produced no effect (Fig. 4c). Together, the results shown in Fig. 4 demonstrate a clear association between the hallucinogenic potential of the compounds, phosphorylation of FOXK2 S360, and lactate production. This finding is in line with recent proteomic studies in human cerebral organoids treated with LSD, which revealed significant alterations in proteins involved in glycolysis and oxidative phosphorylation^31^. However, our phosphoproteomic data reveal for the first time that enhanced glycolytic activity is a distinctive feature of psychedelics with hallucinogenic properties, and that this property is not shared by their non-hallucinogenic analogs. This discovery has potential physiological significance as it links, at the intracellular signaling level, the altered states of consciousness produced by hallucinogenic psychedelics with those induced by physiological anoxia. Indeed, our study suggests that these two interventions converge on enhancing glycolytic metabolism and result in transiently modified states of awareness, which in both cases may ultimately contribute to their restorative or therapeutic effects^32,33^.

**Fig. 4.**
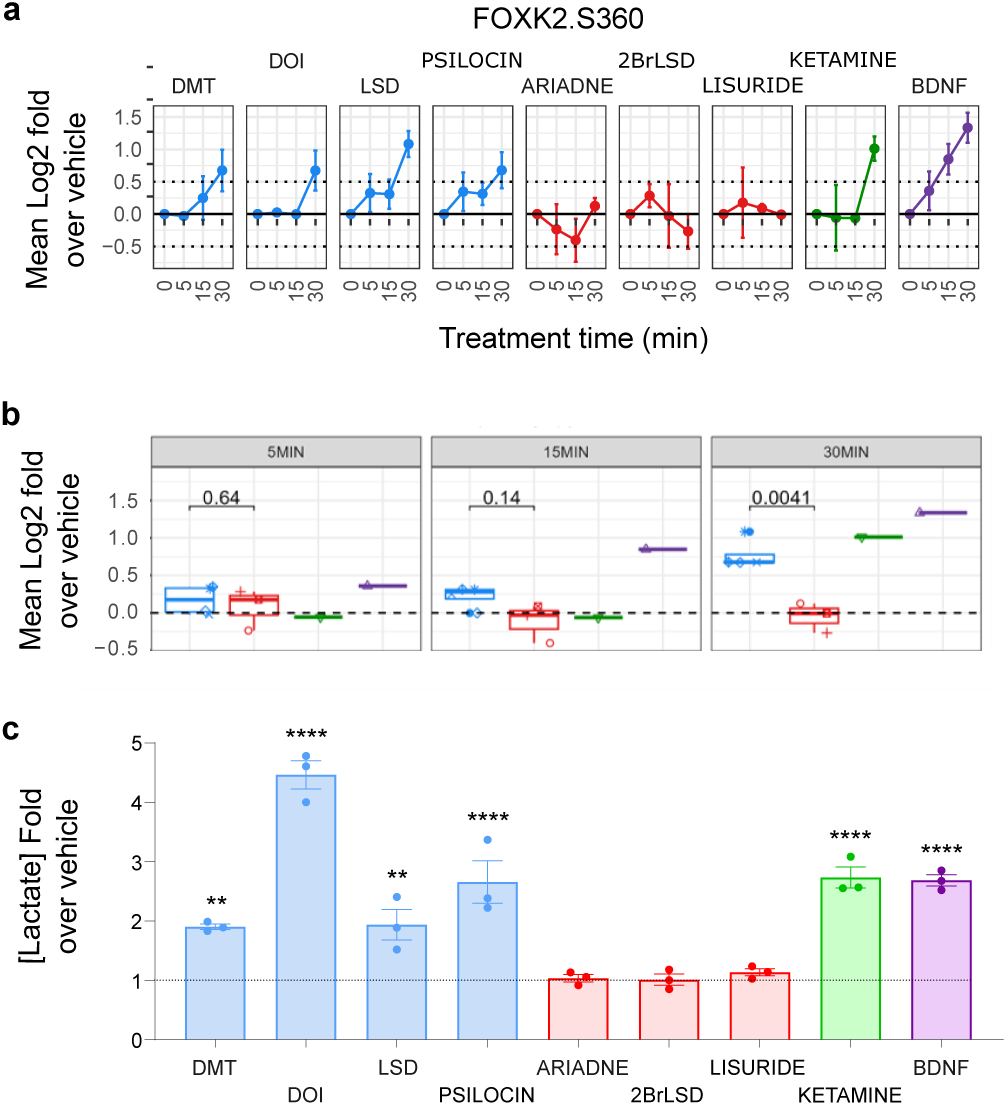
Hallucinogenic psychedelics but not their non-hallucinogenic counterparts induce phosphorylation of FOXK2 at S360 and promote lactate production after treatment. **a.** Phosphorylation of FOXK2 at 5, 15 and 30 minutes post-treatment relativized against vehicle. Datapoints are mean +/- SEM (n = 4 independent replicates) **b.** Area under the curve of the relativized phosphorylation profile of each drug at each timepoint. Statistical differences between hallucinogenic and non-hallucinogenic compounds appear in the 30-minute treatment condition. **c.** Lactate concentration in supernatants from cell cultures treated with the different drugs for 8 hours. Data represent fold change over vehicle. Statistical analysis was performed using one-way repeated measure ANOVA (**C**: F[9,21] = 41.06, p < 0.0001) with Dunnett post-hoc test (*p < 0.05, **p < 0.01, ***p < 0.001, ****p < 0.0001). Data are presented as mean ± SEM of three independent experiments.

In conclusion, this study defines a phosphoproteomic signature of hallucinogenic potential that separates hallucinogenic psychedelics from closely related non-hallucinogenic analogues across multiple chemical classes. By integrating large-scale quantitative phosphoproteomics with a native neural model containing both neurons and glia, we show that psychedelic drugs remodel intracellular signalling networks linking receptor activation to cytoskeletal regulation, chromatin remodelling and cellular metabolism. The resulting map of thousands of regulated phosphorylation events on protein kinases, G-protein–coupled receptors, growth factor receptors and epigenetic enzymes provides a resource for dissecting these pathways in greater detail, while the hallucinogenic signature itself offers a molecular framework for classifying existing compounds and guiding the rational design of new psychedelic-inspired drugs with tailored signalling profiles and potentially dissociable hallucinogenic and neuroplastic effects. Our observation that hallucinogenic psychedelics uniquely engage metabolic modules associated with enhanced glycolytic activity further suggests a convergence with physiological anoxia at the level of cellular energy use, hinting at shared mechanisms that may contribute to altered states of awareness and their restorative or therapeutic impact. Together, these findings establish a conceptual and molecular basis for understanding how distinct psychedelic structures map onto specific intracellular architectures and long-lasting neural adaptations.

## Methods

### Drugs

Brain Derived Neurotrophic Factor (BDNF) was obtained from PreproTech (BDNF, 450-02). Lysergic acid diethylamide (LSD, L7007), (±)-2,5-Dimethoxy-4-iodoamphetamine hydrochloride (DOI, D101) and (±)-ketamine hydrochloride (ketamine, K2753) were purchased from Sigma-Merck. Lisuride maleate was obtained from Tocris (4052) and 2-Bromo-D-lysergic acid diethylamide (2BrLSD, B478840) was purchased from Toronto Research Chemicals (TRC). N,N-Dimethyltryptamine (N,N-DMT, 13959), psilocin (1184) and Ariadne (α-ethyl 2C-D hydrochloride, 26596) were ordered from Cayman Chemical. All drugs were diluted in dimethyl sulfoxide (DMSO) except for BDNF, ketamine and DOI which were diluted in water.

### Cell culture and in vitro drug treatments

Adherent neural stem cell (NSC) lines from the mouse forebrain were obtained following the methodology previously described^12^. Briefly, the fetus dorsal forebrain at E12.5 was dissected and the obtained tissue was disaggregated in order to get a cell suspension. These cells were then transferred to cell-culture plastic plates that had been previously treated with poly-D-lysine 10 μg/ml (Sigma-Merck) to enhance cellular adhesion and promote monolayer cultures. NSCs were maintained and expanded in NS expansion medium composed as follows: DMEM/Ham’s F-12 media with l-glutamine (Gibco), Glucose solution 29mM (Sigma-Merck), MEM nonessential amino acids 1x (Gibco), Penicillin/Streptomycin 1x (Gibco), HEPES buffer solution 4.5 mM (Cytiva), BSA solution 0.012% (Gibco), 2-mercaptoethanol 0.05 mM (Gibco), N2 supplement 1x (Gibco), B27 supplement 1x (Gibco), murine epidermal growth factor (EGF) 10 ng/ml (PeproTech) and human fibroblast growth factor (FGF-2) 10 ng/ml (PeproTech). To induce differentiation of NSCs into various neural cell lineages, the expansion medium was replaced with the following differentiation medium: Neurobasal plus DMEM/Ham’s F-12 media with l-glutamine 1:1 (Gibco), N2 supplement 1x (Gibco), B27 supplement 1x (Gibco), Glutamax 1x (Gibco) and 2-mercaptoethanol 0.05 mM (Gibco). Cells were grown and maintained at 37 °C-5% CO2 under a humidified atmosphere. For in vitro drug treatments, neural stem cells (NSCs) were seeded onto 100 mm culture dishes pre-coated with poly-D-lysine (10 μg/mL) and laminin (3.3 μg/mL). On day five of differentiation, pharmacological agents were added to the culture medium at a final concentration of 10⁻⁵ M, with the exception of Brain Derived Neurotrophic Factor (BDNF), which was administered at 50 ng/mL. Treatments were applied at three distinct time points: 5 minutes, 15 minutes, and 30 minutes. Upon completion of the treatments, cells were harvested and processed for phosphoproteomic analysis. All experiments were independently replicated four times.

### Sample preparation for phosphoproteomics analysis

Mass spectrometry experiments were performed as described previously^34,35^ with some modifications. Cultured cells on 100 mm dishes were washed with ice-cold PBS supplemented with phosphatase inhibitors (1mM Na3VO4 and 1mM NaF). Cells were scrapped and lysed with a Urea buffer (8M urea in 20mM in HEPES, pH 8.0 supplemented with 1mM Na3VO4, 1mM NaF, 1mM Na2H2P2O7 and 1mM sodium β-glycerophosphate) and lysates were then sonicated using a sonicator Vibra-Cells (Sonics Materials) at 20% of intensity, 3 cycles of 10 seconds.

Samples were then centrifuged at 13,000 rpm for 10 min at 5°C. Supernatant was transferred in 1.5mL Protein Lo-bind tube (Eppendorf). Protein concentration was determined using BCA Protein Assay Kit.

For phosphoproteomics analysis, 110 µg of extracted proteins in a volume of 200 ul were reduced with dithiothreitol (DTT, 10 mM) for 1 h at 25 °C, and alkylated with Iodoacetamide (IAM, 16.6 mM) for 30 min at 25 °C. Trypsin beads were equilibrated by three washes with 20 mM HEPES; pH 8.0. Then, samples were diluted with 20 mM HEPES (pH 8.0) to a final concentration of 2 M urea and digested with equilibrated trypsin beads (50% slurry of TLCK-trypsin) overnight at 37 °C. Trypsin beads were subsequently removed by centrifugation (2000×g for 5 min at 5 °C) and samples were transferred to 96 well plates and acidified by adding TFA to a final concentration of 0.1%. Peptide solutions were desalted and subjected to phosphoenrichment using the AssayMAP Bravo (Agilent Technologies) platform. For desalting, protocol peptide clean-up v3.0 was used. Reverse phase S cartridges (Agilent, 5 μL bed volume) were primed with 250 μL 99.9% acetonitrile (ACN) with 0.1%TFA and equilibrated with 250 μL of 0.1% TFA at a flow rate of 10 μL/min. The samples were loaded (770 μL) at 20 μL/min, followed by an internal cartridge wash with 250 μL of 0.1% TFA at a flow rate of 10 μL/min.

Peptides were then eluted with 105 μL of 1M glycolic acid with 50% ACN, 5% TFA and this is the same buffer for subsequent phosphopeptide enrichment.

For phospho-enrichment, we followed the Phospho Enrichment v 2.1 protocol on the Assay MAP Bravo platform with the following modifications. The 5µl Assay MAP TiO2 cartridges were primed with 100µl of 5% ammonia solution with 15% ACN at a flow rate of 300 μL/min and equilibrated with 50 μL loading buffer (1M glycolic acid with 80% ACN, 5% TFA) at 10 μL/min. Samples eluted from the desalting were loaded onto the cartridge at 3 μL/min. The cartridges were washed with 50 μL loading buffer and the phosphorylated peptides were eluted with 25 μL 5% ammonia solution with 15% ACN directly into 25 μL 10% formic acid.

Phosphopeptides were lyophilized in a vacuum concentrator and stored at -80°C. For proteomics analysis, 30 μg of protein were digested and acidified as described above. Peptide solutions were desalted using the AssayMAP Bravo (Agilent Technologies) platform. Cartridges were primed, equilibrated, loaded and washed as described above and peptides were eluted with 105 μL of 70/30 ACN/ H2O + 0.1% TFA. Eluted peptide solutions were dried in a SpeedVac vacuum concentrator and peptide pellets were stored at −80 °C.

### LC-MS/MS Analysis

Peptides were re-suspended in 20 µL of reconstitution (97% H2O, 3% ACN, 0.1% TFA, 50fmol/µl-1 enolase peptide digest) and sonicated for 5 minutes at RT. Following a brief centrifugation, 2μl was loaded onto a LC-MS/MS system. This consisted of a nano flow ultra-high pressure liquid chromatography system UltiMate 3000 RSLC nano (Dionex) coupled to a Q Exactive Plus using an EASY-Spray system. The LC system used mobile phases A (3% ACN: 0.1% FA) and B (100% ACN; 0.1% FA). Peptides were loaded onto a μ-pre-column and separated in an analytical column. The gradient: 1% B for 5 min, 1% B to 35% B over 90min, following elution the column was washed with 85% B for 7 min, and equilibrated with 3% B for 7min, flow rate of 0.25 µL/min. Peptides were nebulized into the online connected Q-Exactive Plus system operating with a 2.1s duty cycle. Acquisition of full scan survey spectra (m/z 375-1,500) with a 70,000 FWHM resolution was followed by data-dependent acquisition in which the 15 most intense ions were selected for HCD (higher energy collisional dissociation) and MS/MS scanning (200-2,000 m/z) with a resolution of 17,500 FWHM. A 30s dynamic exclusion period was enabled with an exclusion list with 10ppm mass window. Overall duty cycle generated chromatographic peaks of approximately 30s at the base, which allowed the construction of extracted ion chromatograms (XICs) with at least ten data points.

### Peptide and protein identification and quantification

Peptide identification from MS data was automated using a Mascot Daemon workflow in which Mascot Distiller generated peak list files (MGF) from RAW data, and the Mascot search engine matched the MS/MS data stored in the MGF files to peptides using the SwissProt Database restricted to Mus musculus (SwissProt_2024_03.fasta). Searches had an FDR of ∼1% and allowed 2 trypsin missed cleavages, mass tolerance of ±10 ppm for the MS scans and ±25 mmu for the MS/MS scans, carbamidomethyl Cys as a fixed modification and oxidation of Met, PyroGlu on N-terminal Gln and phosphorylation on Ser, Thr, and Tyr as variable modifications. Percolator was applied to improve the discrimination between correct and incorrect spectrum identifications^36^. Pescal was used for label free quantification of the identified peptides^37^. The software constructed XICs for all the peptides identified in at least one of the LC-MS/MS runs across all samples. XIC mass and retention time windows were ±7 ppm and ±2 min, respectively. Quantification of peptides was achieved by measuring the area under the peak of the XICs.

Individual peptide intensity values in each sample were normalized to the sum of the intensity values of all the peptides quantified in that sample. Phosphoproteomics data was processed and analysed using a bioinformatic pipeline developed in a R environment (https://github.com/CutillasLab/protools2/). The normalised data was centred, log2 scaled and 0 values were inputted using the minimum feature value in the sample divided by five, considering the log2 scale. Then, p-values to assess statistical differences between comparisons were calculated using LIMMA and then adjusted for FDR using the Benjamini-Hochberg procedure. The mass spectrometry proteomics data have been deposited to the ProteomeXchange Consortium via the PRIDE^38^ partner repository with the dataset identifier PXD070037 and 10.6019/PXD070037.

### Kinase substrate enrichment analysis, KSEA

Kinase activity was inferred from the phosphoproteomics dataset using kinase substrate enrichment analysis (KSEA)^39^, available in our bioinformatic pipeline *protools2*. In summary, this approach evaluates kinase activity by measuring the phosphorylation levels of known substrate proteins associated with each kinase.

### Lactate measurements

Lactate release into the culture medium following drug treatments was quantified using the L-Lactate Assay Kit (Sigma–Merck, MAK329) according to the manufacturer’s instructions.

Neural stem cells (NSCs) were seeded onto 12-well plates coated with poly-D-lysine (10 μg/mL) and laminin (3.3 μg/mL). On day five of differentiation, cells were treated with the respective compounds under conditions equivalent to those used for the phosphoproteomics experiments.

One hour prior to drug administration, the culture medium was replaced with fresh medium. Culture medium was collected on ice at 1, 2, 4, and 8 hours post-treatment, centrifuged for 10 minutes at 2,500 rpm at 4 °C, and stored at –80 °C until analysis to determine the optimal time point for sample collection in later experiments. For lactate quantification, samples were collected 8 hours after drug treatment. Lactate levels were normalized to total cellular protein content.

### Head Twitch Response (HTR) assays

DMT (*N*,*N*-dimethyltryptamine) was obtained from Sigma (SML0791) and dissolved at 25 mg/ml in 100% DMSO, then diluted to 0.6 mg/ml on the day of the experiment in sterile 0.9% saline to achieve a 3 mg/kg dose via 5 µl/g injection volume. DOI [(±)-1-(2,5-dimethoxy-4-iodophenyl)-2-aminopropane] was obtained from Sigma (D101) and dissolved in sterile 0.9% saline to 0.4 mg/ml to achieve a 2 mg/kg dose via 5 µl/g injection volume. LSD (lysergic acid diethylamide) was obtained from Lipomed (397-FB) and was dissolved at 10 mg/ml in 100% DMSO, then diluted to 0.04 µg/ml on the day of the experiment in sterile 0.9% saline to achieve a 0.2 mg/kg dose via 5 µl/g injection volume (i.p). Psilocybin was obtained from Usona (non-cGMP) and was dissolved in sterile 0.9% saline to 0.2 mg/ml to achieve a 1 mg/kg dose via 5 µl/g injection volume (i.p). Ariadne (1-(2,5-dimethoxy-4-methylphenyl)butan-2-amine) was obtained from Cayman Chemicals (26596) and was dissolved in sterile 0.9% saline to 0.6 mg/ml to achieve a 3 mg/kg dose via 5 µg/ml injection volume (i.p). 2BrLSD was obtained from Toronto Research Chemicals (TRC-B478840) and was dissolved to 14 mg/ml in 100% DMSO, then diluted in sterile 0.9% saline to 0.6 mg/ml to achieve a dose of 3 mg/kg dose via 5 µl/g injection volume (i.p). Lisuride maleate was obtained from Tocris (4052) and dissolved to 2 mg/ml in 100% DMSO, then was diluted in sterile 0.9% saline to 0.08 mg/ml to achieve a dose of 0.4 mg/kg via 5 µl/g injection volume (i.p). Ketamine HCl Injectable Solution (100 mg/ml) was obtained from Covetrus (71069) and diluted to 0.6 mg/ml to achieve a dose of 3 mg/kg dose via 5 µl/g injection volume (i.p). 7,8-Dihydroxyflavone (DHF) was obtained from Abcam (AB120996) and dissolved in 100% DMSO, then was diluted in sterile 0.9% saline to 1 mg/ml to achieve a dose of 5 mg/kg via 5 µl/g injection volume (i.p). The vehicle groups consisted of 3 mice that received sterile 0.9% saline (5 µl/g, i.p.) and 3 mice that received 30% DMSO dissolved in sterile 0.9% saline (5 µl/g, i.p.) to account for the maximum DMSO concentration represented in the DMSO-dissolved compounds. These two vehicle groups were not statistically significant from each other in the HTR assay (data not shown), and were therefore combined into a single “vehicle control” group. Doses were selected based on prior studies demonstrating behavioral effect of the tested drugs^6,9,10,40,41^. Experiments were performed on adult (8-10 weeks of age) male C57BL/6J mice (The Jackson Laboratory, Ban Harbork, ME). Animals were housed in cages with up to 4 littermates at 12 h light/dark cycle at 23 °C with food and water ad libitum, except during behavioral testing. Experiments were conducted in accordance with NIH guidelines, and were approved by the Virginia Commonwealth University Animal Care and Use Committee. All efforts were made to minimize animal suffering and the number of animals used. Detection of head-twitch responses (HTR) in mice was performed as previously reported^42,43^.

Briefly, mice were ear-tagged with neodymium magnets (N50, 3 mm diameter × 1 mm height, 50 mg) glued to the top surface of aluminum ear tags for rodents (Las Pias Ear Tag, Stoelting Co.) with the magnetic south of the magnet in contact with the tag. Following ear-tagging animals were placed back into their home cages and allowed to become accustomed to the tags for one week. Data acquisition and data processing was performed as previously described^42,43^, using our signal analysis protocol in combination with a deep learning-based protocol based on scalograms. Testing occurred no more than once per week with at least 7 days between test sessions. On test days, mice were placed individually into the monitoring chamber for 15 min to acclimate to the environment and determine baseline HTR. Subsequently, the animals received the corresponding treatments and HTRs were recorded for 90 minutes.

### Statistics

In phosphoproteomics studies statistical analysis was carried out in R (v4.3.1) using the limma package, base functions or using the ggpubr package. Gene Ontology analysis of genes/proteins differentially was performed using functions in the package clusterProfiler or those in protools2. Data was visualized using the ggplot2 package. In HTR assays animals were randomly allocated into the different experimental groups. No statistical methods were used to predetermine sample sizes, but our sample sizes are similar to those reported in previous publications from our and other laboratories. Statistical significance was assessed by one-way or two-way repeated measures ANOVA, depending upon the number of experimental conditions and independent variables. Following a significant ANOVA, specific comparisons were made using Bonferroni’s or Fisher’s LSD post-hoc tests. All statistical analyses were performed with GraphPad Prism software version 10, and comparisons were considered statistically significant if p < 0.05. All values in the figures represent mean ± S.E.M.

## Acknowledgments

We thank Vinothini Rajeeve and Tommy Shields (CRUK Barts Centre Mass Spectromtry Facility) for their support for their guidance and technical assistance with the proteomics experiments. This research was supported by the Spanish Ministry of Science, Innovation and Universities (MCIN) and the State Research Agency (AEI, grant PID2021-125448OB-I00), co-funded by the European Regional Development Fund (FEDER, “A way of making Europe”) (JFLG). NIH R01MH084894 (JGM) and NIH T32DA007027 (JLM). CRUK (C28206/A14499 and C15966/A24375), and Medical Research Council (MC_PC_ MR/X013766/1) (PRC).

## Author contributions

Conceptualization: JFLG, JGM, PRC. Methodology: SMMG, MTT, JLM, SP, JGM, PRC, JFLG. Investigation: SMMG, MTT, JLM, SP, JGM, PRC, JFLG. Funding acquisition: JFLG, JGM, PRC. Supervision: JFLG, JGM, PRC. Writing – original draft: JFLG, PRC. Writing – review & editing: JFLG, JGM, PRC.

## Data and materials availability

The mass spectrometry proteomics data have been deposited to the ProteomeXchange Consortium via the PRIDE partner repository with the dataset identifier PXD070037 and 10.6019/PXD070037. Phosphoproteomics and proteomics data was processed and analyzed using a bioinformatic pipeline developed in a R environment (https://github.com/CutillasLab/phosphopsychedelics/).

---

**Extended Data Fig.1.**
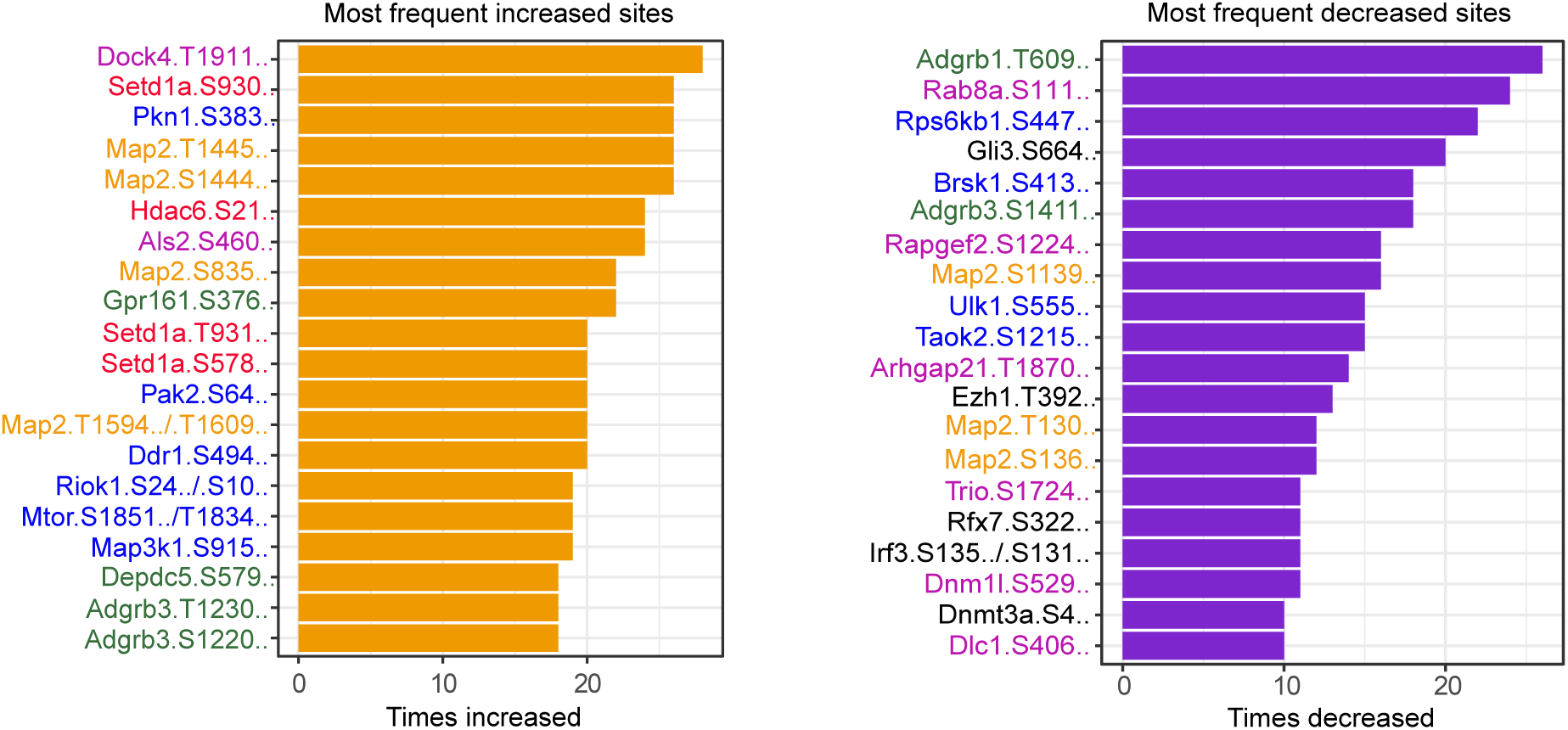
Most frequently increased and decreased phosphorylation sites induced by psychedelic compounds and BDNF are of on proteins with functions on kinase and GPCR signaling and on epigenetic regulators. The most frequently increased phosphorylation sites across treatments were observed on several protein kinases (blue fonts), including the Ser/Thr kinases Pkn1 at S383 (a PKC family member), Pak2 at S64, Riok1 at S24, Mtor at S185, and Map3k1 at S915, as well as on the receptor tyrosine kinase Ddr1 at S494. Additional increase in phosphorylation were detected on GTPase regulators such as Als2 and Dock4 (purple), the GPCRs Gpr161 and Adgrg3 (green), the epigenetic regulators Setd1a (at three distinct sites) and Hdac6 (red in Fig. 1F), and three sites on Map2 (orange), a cytoskeletal protein with a well-established role in neuronal development. Conversely, the most frequently decreased sites were found on the GPCRs Adgrb1 and Adgrb3; small GTPases and GTPase regulators including Dnm1l, Rab8a, Arhgap21, Rapgef2, Trio, and Dlc1; kinases such as Rps6kb1, Brsk1, Ulk1, and Taok2; and transcriptional activators including Gli3, Irf3, and Rfx7.

**Extended Data Fig. 2.**
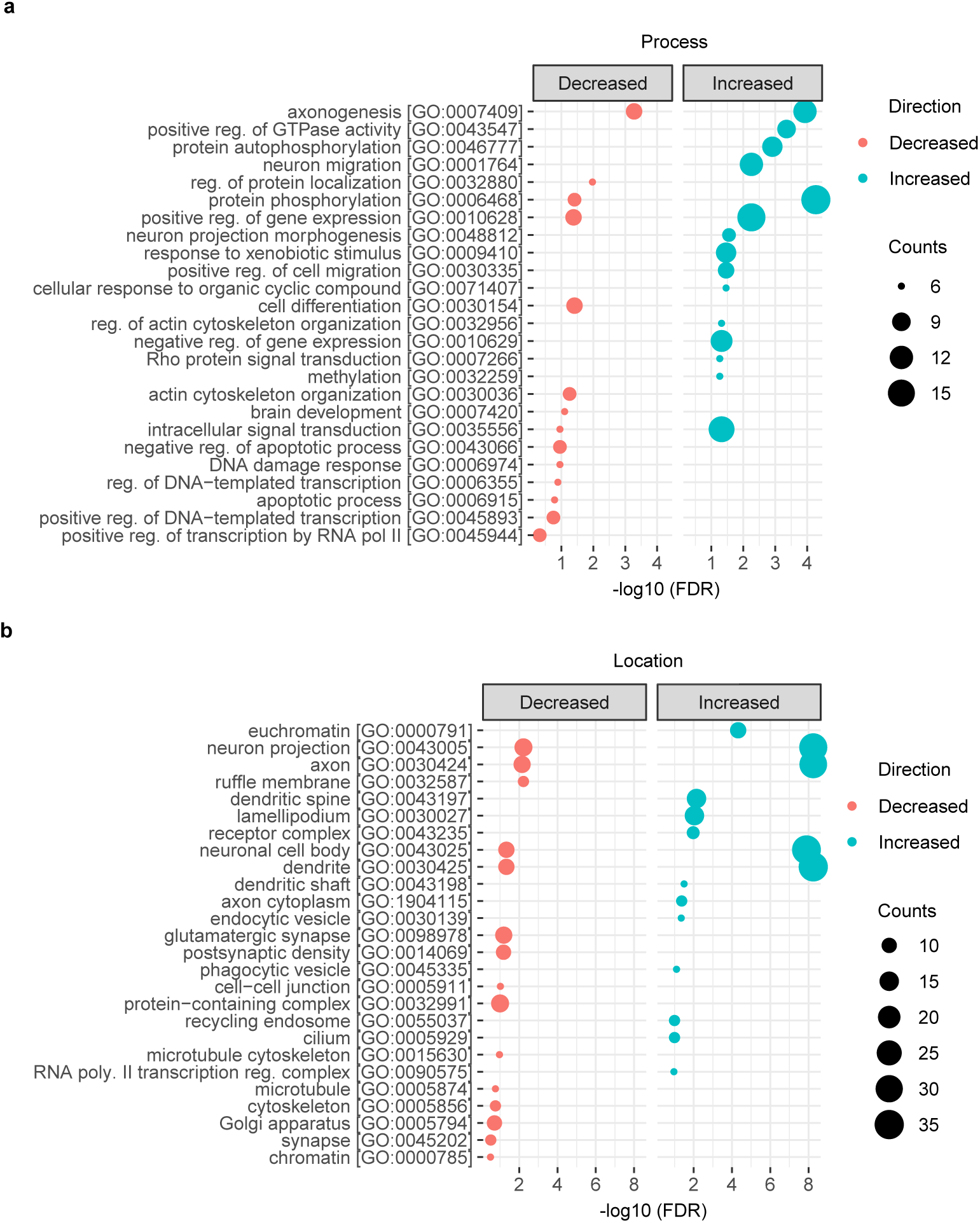
Ontology enrichment analysis of regulated phosphoproteins. **a.** Biological processes, and **b.** Cellular component after treatment with BDNF and psychedelic drugs.

**Extended Data Fig. 3.**
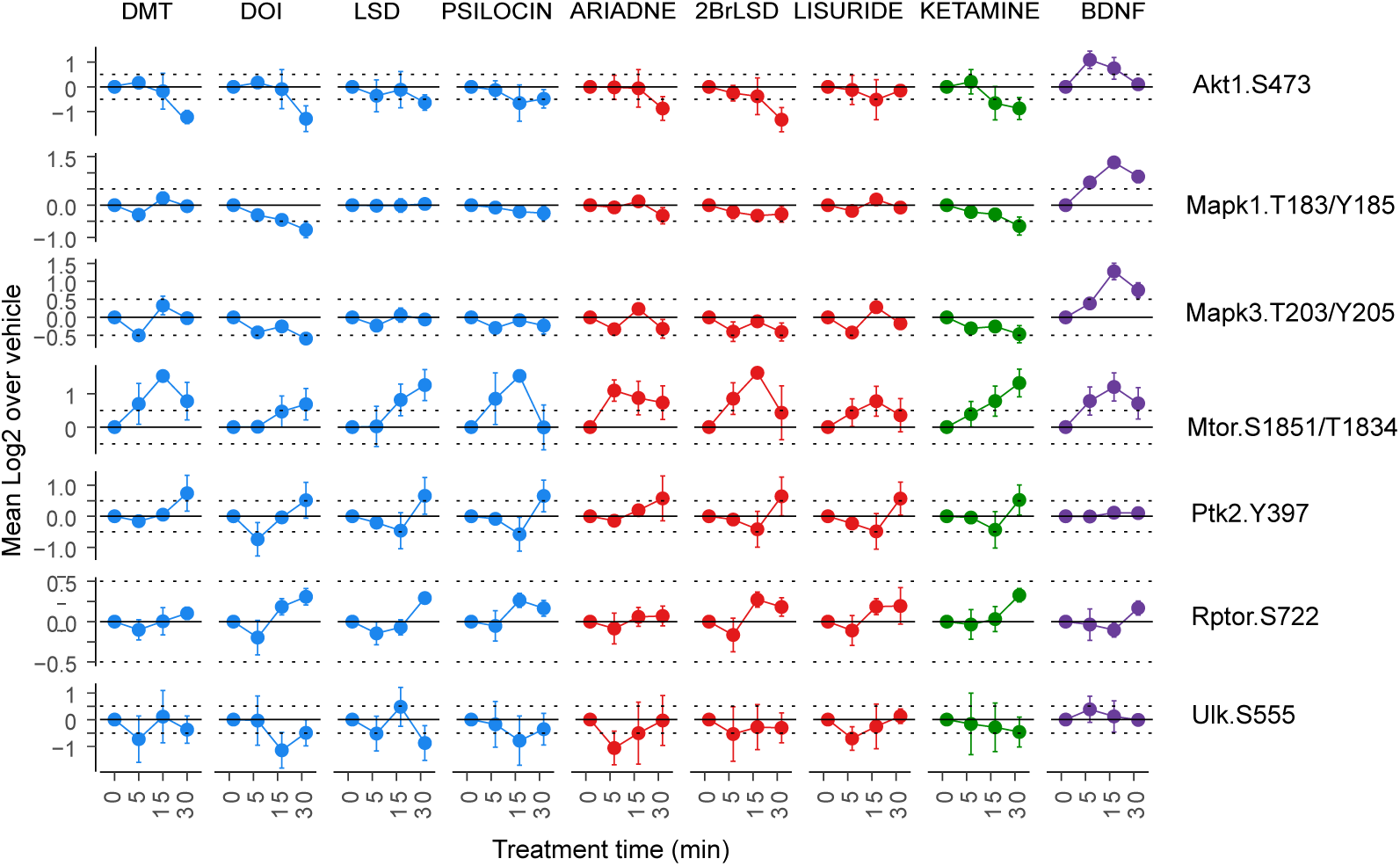
Examples of phosphopeptides linked to kinase activities identified by KSEA analysis. To validate the findings obtained through KSEA, we examined the phosphorylation status of canonical kinase activity markers within the AKT and MAPK pathways. Phosphorylation of PKB (Akt1) at S473, an activation site on this kinase, was decreased by all psychedelic treatments but increased following BDNF exposure. Similarly, activating phosphorylation sites on ERK1 and ERK2 (genes Mapk3 and Mapk1, respectively) increased in response to BDNF but decreased or remained unchanged after treatment with psychedelics. In contrast, phosphorylation of mTOR at S1851 markedly increased under all treatment conditions, whereas phosphorylation of the AMPK substrate ULK1 tended to decrease after exposure to psychedelics and remained unaltered in cells treated with BDNF. We also observed increased phosphorylation at the activating site Y397 of focal adhesion kinase FAK1 (Ptk2) following 30-minute treatments with all psychedelics, but not with BDNF. This kinase regulates cellular migration, adhesion, and spreading, and its activation pattern after exposure to psychedelics is consistent with the enrichment of neuron migration ontologies observed previously (Extended Data Fig. 2).

**Extended Data Fig. 4.**
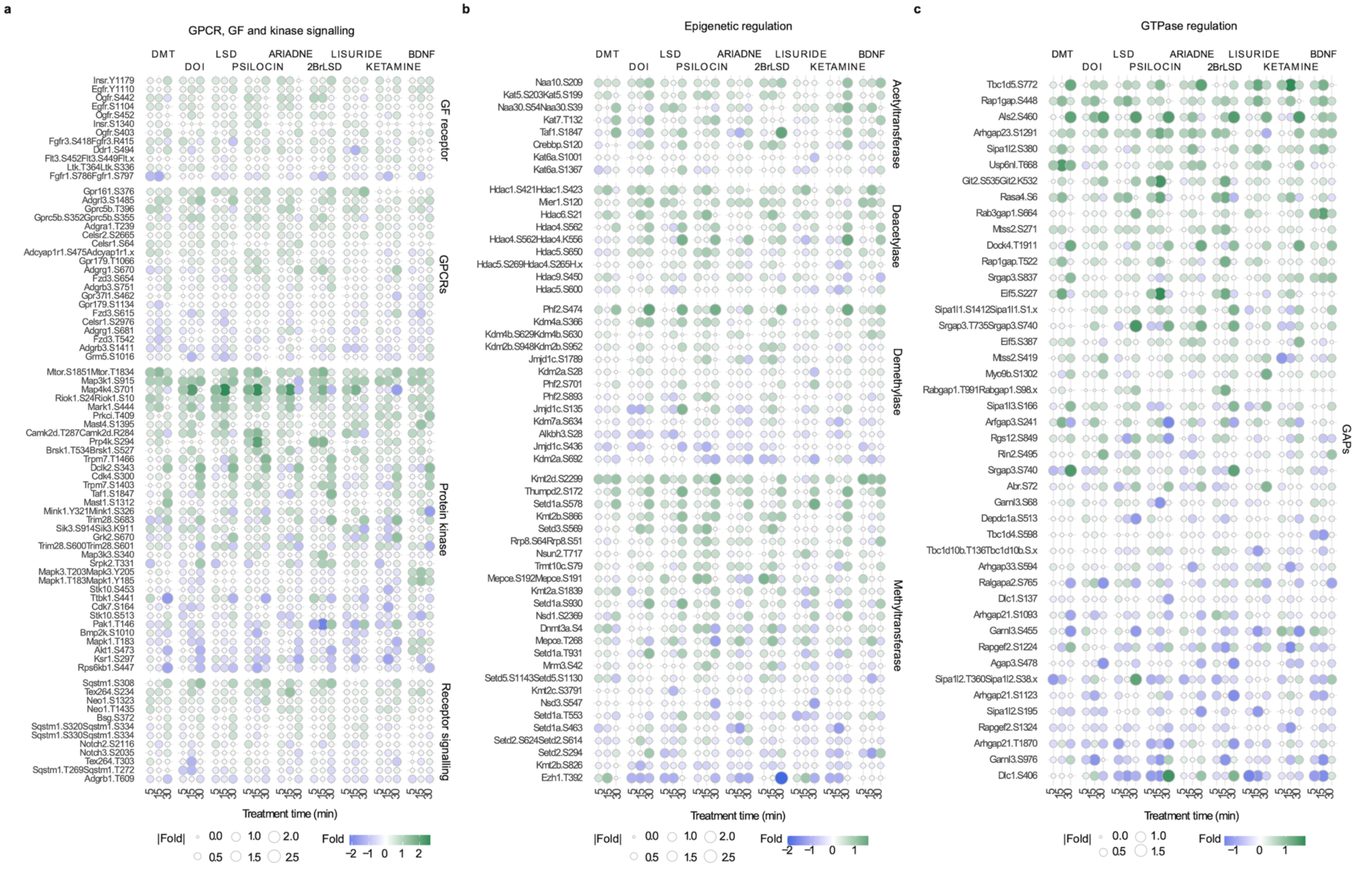
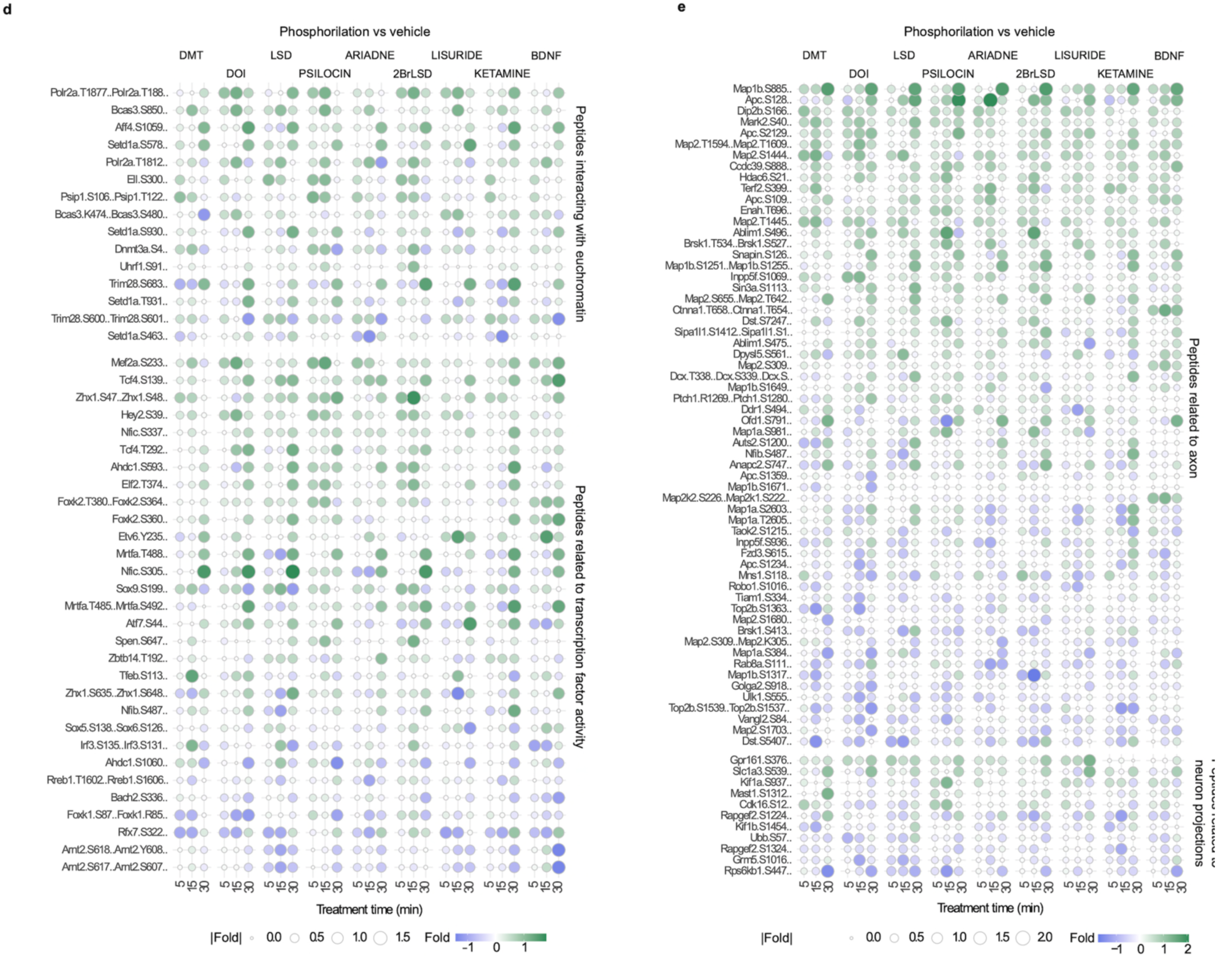
Effect of BDNF and psychedelic drugs in the phosphorylation of proteins linked to different signalling pathways. Protein phosphorylation of proteins linked to multiple growth factor receptors (GFRs) and GPCRs signalling (**a**), epigenetic regulation (**b**), GTPase regulation (**c**), euchromatin and transcription regulation (**d**), and axon and neuron projection (**e**). Datapoints show the Log2 Fold for each phosphopeptide across treatments and treatment times, with size and color representing the values of Log2 Fold differences between treatment and control at the different time points.

**Extended Data Fig. 5.**
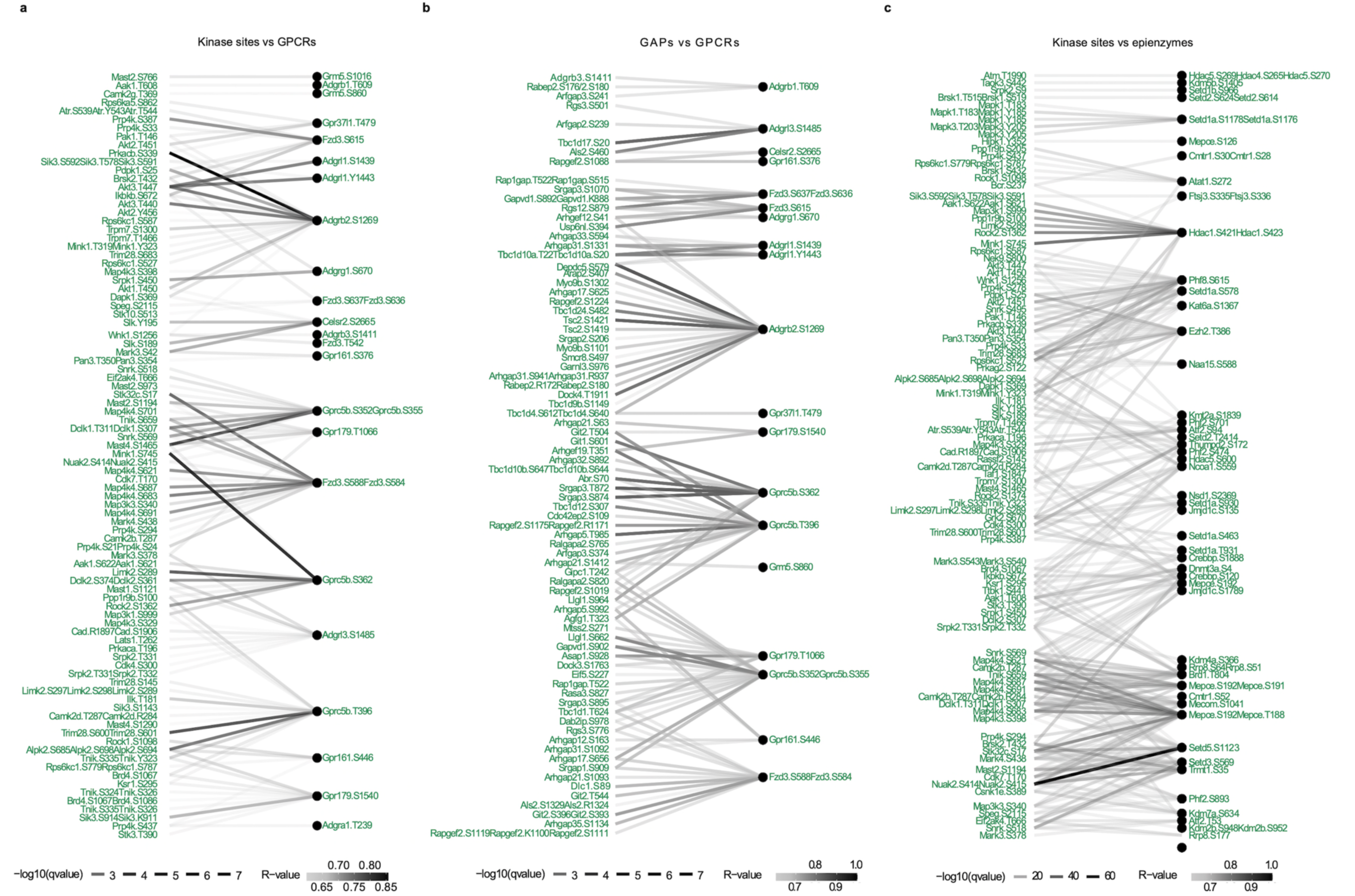
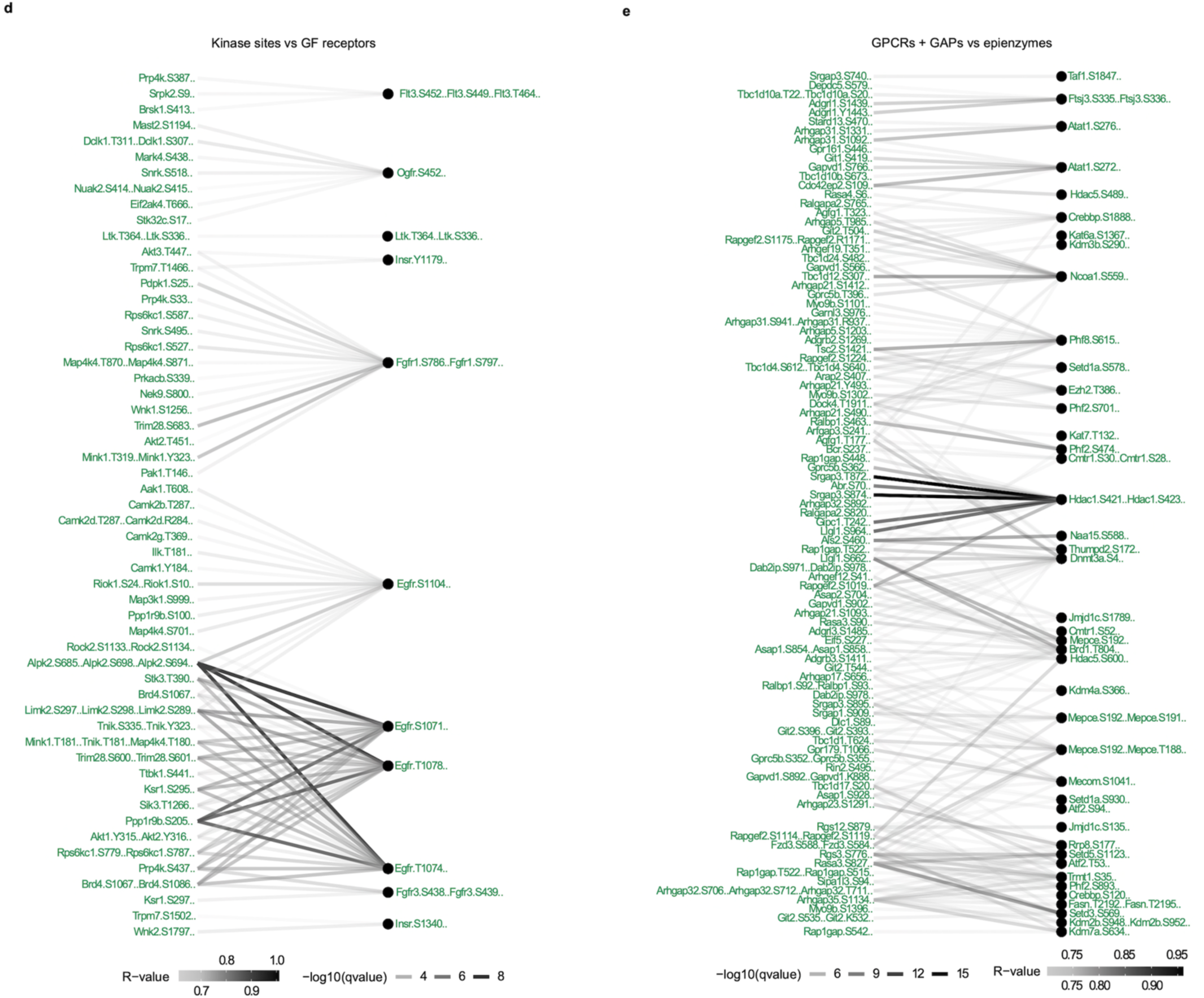
Phosphorylation regulatory network depicting associations between different phosphosites across treatments. Nodes represent phosphosites and the edges represent the correlation coefficients between the temporal phosphorylation of the named nodes by the different drugs. **a, b, c, d, e**. Phosphorylation networks showing the relationship between phosphosites on kinase sites and on GPCRs (a), GTPase-activating proteins (GAPs) and GPCRs (b), kinase sites and epigenetic regulators (c). kinase sites and grow factor (GF) receptors (d), and GTPase-activating proteins (GAPs)/GPCRs and epigenetic regulators (e)

**Extended Data Fig. 6.**
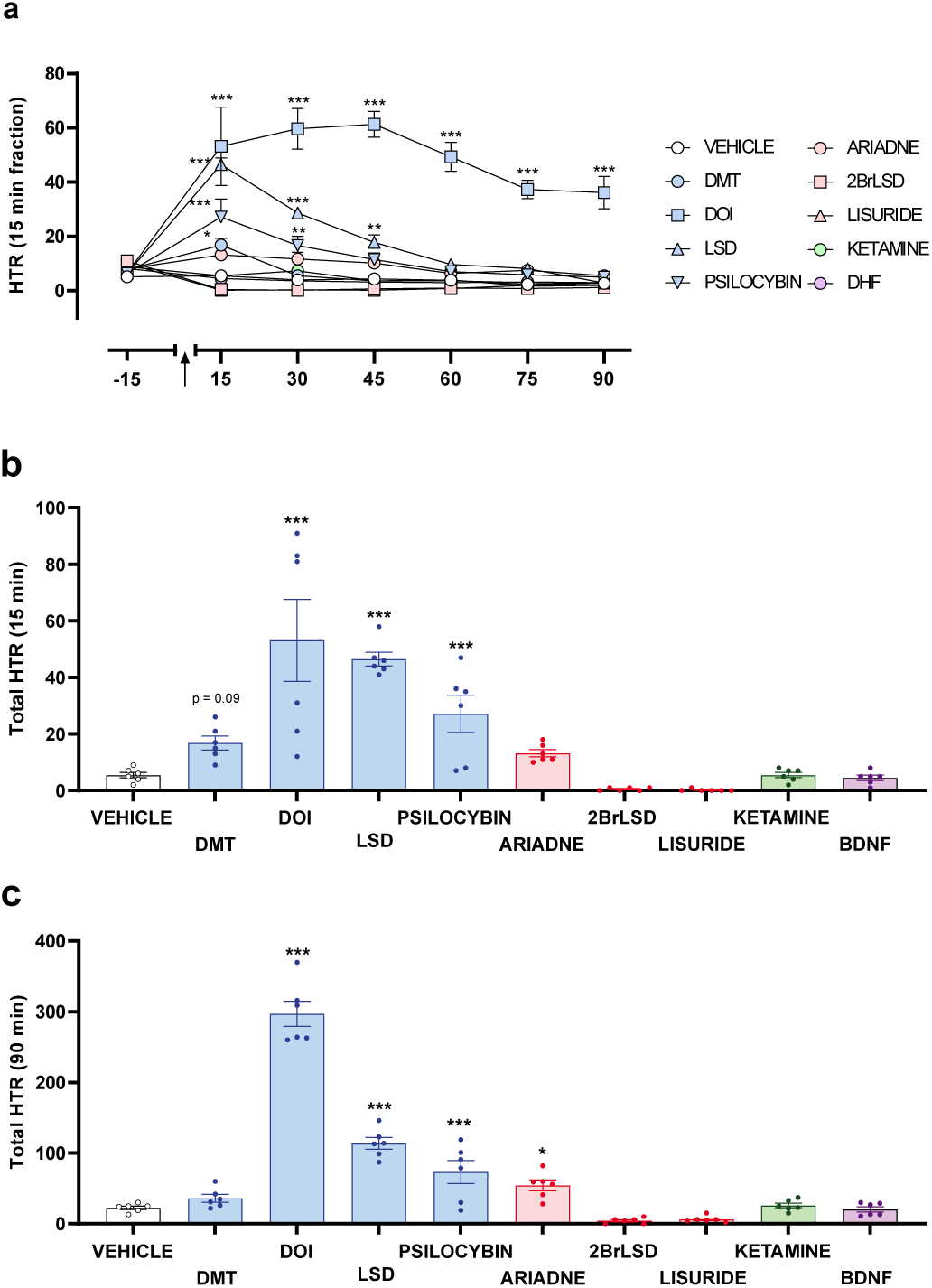
Head Twitch Response (HTR) assays. Effect of DMT (3 mg/kg), DOI (2 mg/kg), LSD (0.2 mg/kg), psilocybin (1 mg/kg), Ariadne (3 mg/kg), 2BrLSD (3 mg/kg), lisuride (0.4 mg/kg), ketamine (3 mg/kg) and DHF (5 mg/kg), or vehicle administration (i.p.) on HTR in mice (n = 6 mice per group). **a.** Time-course showing HTR counts in 15-min blocks corresponding to different drugs or vehicle. Black arrow shows the administration time-point of drug or vehicle. **b.** HTR counts correspond to the first 15 min after drug or vehicle administration. **c.** HTR counts correspond to the first 90 min after drug or vehicle administration. Statistical analysis was performed using two-way repeated measure ANOVA (A: drug F[9,50] = 90.97, p < 0.001; time F[6,300] = 24.12, p < 0.001; interaction F[54,300] = 9.75) with Bonferroni post-hoc test and one-way ANOVA (B: F[9,50] = 13.42, p < 0.001; C: F[9,50] = 99.91 p < 0.001) with Fisher’s LSD post-hoc (*p < 0.05, **p < 0.01, ***p < 0.001). Data are presented as mean ± SEM.

**Extended Data Fig. 7.**
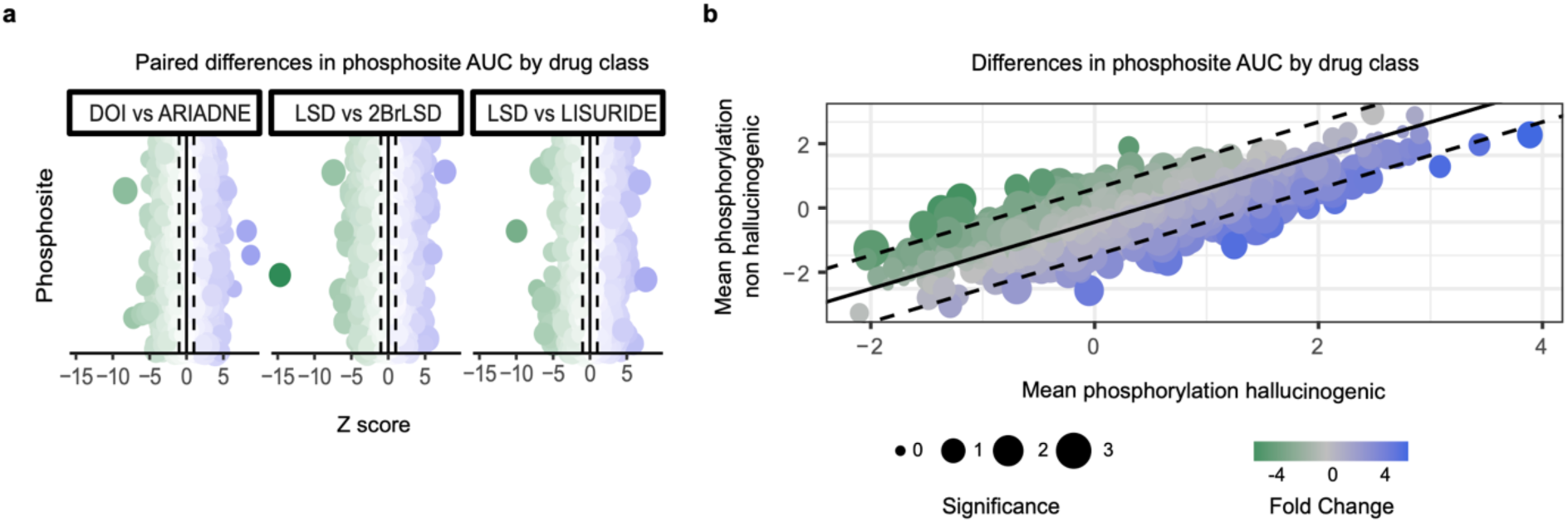
Approach to identify a phospho-signature of psychedelic activity. **a.** Areas under the curve (AUC), calculated for each phosphopeptide, were compared across hallucinogenic compounds and their non-hallucinogenic structural homologues. The dots represent the differences, expressed as Z score, for each phosphopeptide. **b.** Scatter plot showing the mean phosphorylation of phosphopeptides induced by different classes of psychedelics. Phosphosites consistently increased or decreases by hallucinogenic compounds relative to the effects of non-hallucinogenic drugs were included in the signature.

**Extended Data Fig. 8.**
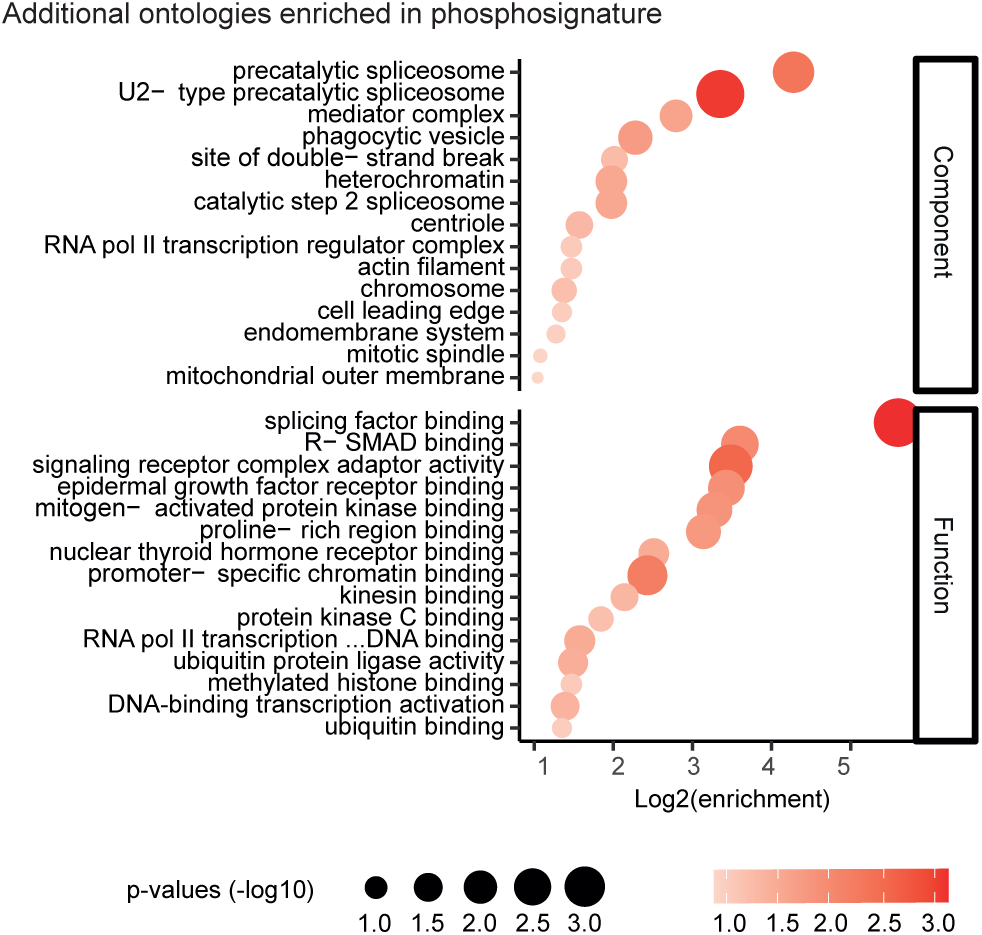
Gene Ontology analysis corresponding to differential phosphopeptides between hallucinogenic versus non-hallucinogenic psychedelic drugs. The color and size of the dots show the significance of each term. The x-axis shows the enrichment of each gene ontology term.

**Supplementary Fig. 1.**
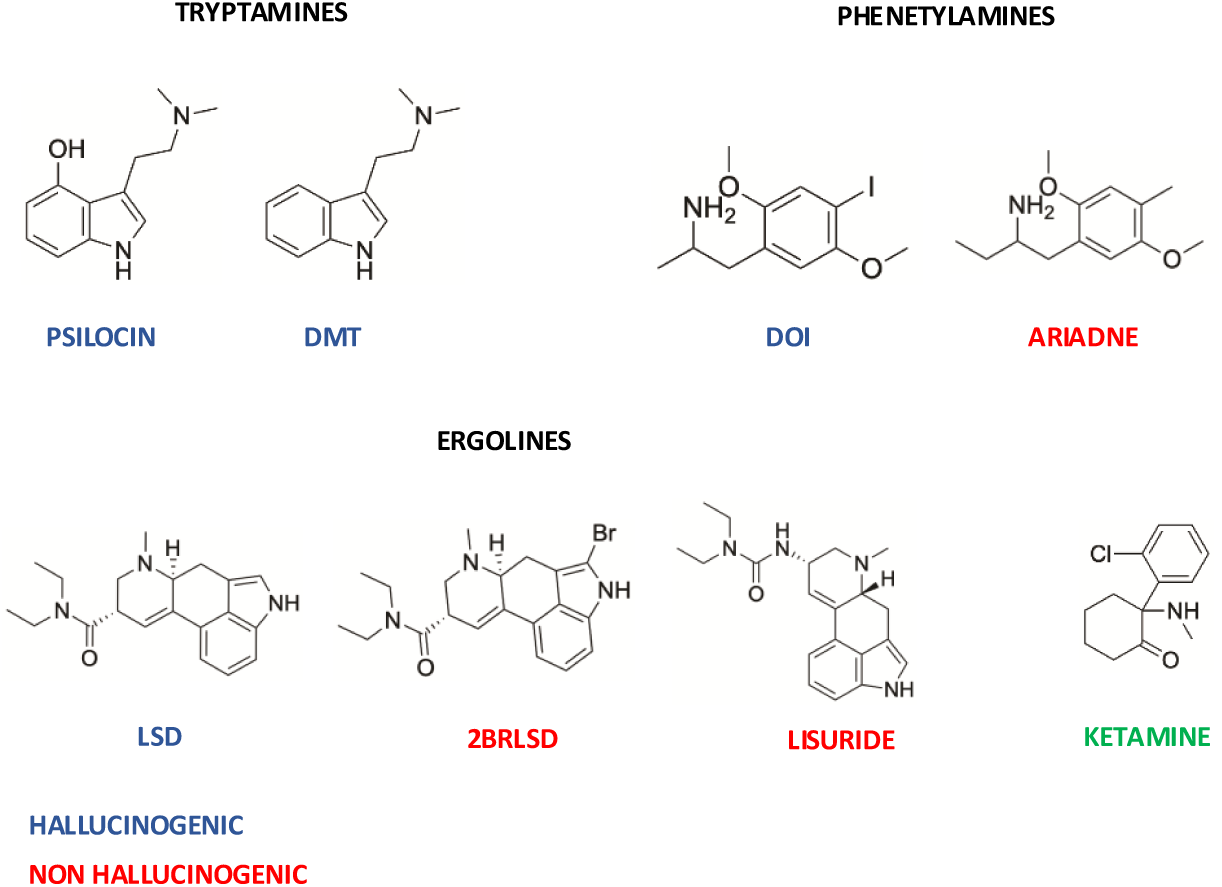
Chemical structures of psychedelic drugs used in this study. Serotonergic psychedelics are grouped according to their chemical class, tryptamines, phenethylamines, and ergolines. Hallucinogenic and non-hallucinogenic compounds are indicated in blue and red, respectively. Structures were drawn using ChemDraw.

